# YfmR is a translation factor that prevents ribosome stalling and cell death in the absence of EF-P

**DOI:** 10.1101/2023.08.04.552005

**Authors:** Hye-Rim Hong, Cassidy R. Prince, Daniel D. Tetreault, Letian Wu, Heather A. Feaga

## Abstract

Protein synthesis is performed by the ribosome and a host of highly conserved elongation factors. Elongation factor P (EF-P) prevents ribosome stalling at difficult-to-translate sequences, particularly polyproline tracts. In bacteria, phenotypes associated with *efp* deletion range from modest to lethal, suggesting that some species encode an additional translation factor that has similar function to EF-P. Here we identify YfmR as a translation factor that is essential in the absence of EF-P in *B. subtilis*. YfmR is an ABCF ATPase that is closely related to both Uup and EttA, ABCFs that bind the ribosomal E-site and are conserved in more than 50% of bacterial genomes. We show that YfmR associates with actively translating ribosomes and that depleting YfmR from Δ*efp* cells causes severe ribosome stalling at a polyproline tract *in vivo*. YfmR depletion from Δ*efp* cells was lethal, and caused reduced levels of actively translating ribosomes. Our results therefore identify YfmR as an important translation factor that is essential in *B. subtilis* in the absence of EF-P.

**Significance:** Translation is one of the most ancient and energetically demanding processes that occurs in the cell. Ribosomes constitute more than 60% of cellular mass in actively growing cells, and ribosomes are a major target of antimicrobials and chemotherapeutics. Here, we identify YfmR as a translation factor that is essential in the absence of EF-P. YfmR is a member of the ABCF family of ATPases whose role in translation is only beginning to be understood. Given the broad distribution of ABCFs from bacteria to fungi, we expect our results to have implications for understanding translation elongation in diverse organisms.

## Introduction

Ribosomes catalyze peptide bond formation by using a series of tRNAs to decode an mRNA template. In bacteria, translation initiates when an mRNA and the initiation factors IF1, IF2, and IF3 bind the small 30S ribosomal subunit (1). IF2 and IF3 ensure that a specialized tRNA charged with formyl-methionine (tRNA^fMet^) is positioned at the P-site during initiation (2–6). After IF3 dissociates, the large 50S subunit joins to form the 70S initiation complex (70SIC) (7). IF1 and IF2 then dissociate, and charged tRNAs are delivered to the A-site of the 70S elongation complex (70SEC) by EF-Tu (8, 9). The large ribosomal subunit catalyzes peptide bond formation and the nascent chain is transferred from the P-site tRNA to the A-site tRNA (10–13). The ribosome is then advanced one codon by EF-G (14–16). Elongation proceeds at a rate of approximately 15 codons sec^-1^ (17) and terminates at a stop codon.

Ribosomes become stalled at polyproline tracts because the cyclical nature of proline causes an unfavorable conformation of the nascent peptide in the exit tunnel that destabilizes the peptidyl-tRNA in the P-site (18). In bacteria, the near universally conserved elongation factor EF-P prevents ribosome stalling at prolines by stabilizing the P-site tRNA and forcing the nascent chain to adopt a favorable geometry for peptide bond formation (18–25). Structural and biochemical data also support a role for EF-P in promoting formation of the first peptide bond (26–28). EF-P stimulates peptide bond formation *in vitro* between fMet-Lys, fMet-Leu, and fMet-Gly, though not between fMet-Phe (26, 29). Both tRNA^fMet^ and tRNA^Pro^ are unique in that they contain a C17pU17a bulge in the D-loop, which could serve as a specificity determinant for EF-P (18, 30–32).

EF-P is essential in some bacterial species, such as *Acinetobacter baumannii* (33), *Neisseria meningitidis* (34), and *Mycobacterium tuberculosis* (35), but is dispensable in others, such as *Salmonella enterica* (36), *Escherichia coli* (37–39), and *Bacillus subtilis* (37, 40–44). Since *efp* deletion phenotypes range from modest to lethal in different bacterial species, we hypothesized that some bacteria encode an uncharacterized translation factor with a similar function. To identify this factor, we used transposon mutagenesis coupled to Illumina sequencing (Tn-seq) to screen for genes that are essential in the absence of EF-P in *B. subtilis*. This screen identified *yfmR* as a gene that is essential when *efp* is deleted. We find that depleting YfmR from Δ*efp* cells causes severe ribosome stalling at a polyproline tract and decreases actively translating ribosomes. We also find that deleting *efp* from *Bacillus anthracis* causes severe growth and sporulation defects and that heterologous expression of *B. subtilis* YfmR in *B. anthracis* Δ*efp* cells partially rescues these phenotypes. These results suggest that YfmR and EF-P have similar functions in preventing ribosome stalling, and that this function is essential in *B. subtilis*.

## Results

### Tn-seq identifies genetic interactions with *efp*

EF-P is essential in some bacterial species but dispensable in others, suggesting that some species encode translation factors with compensatory or redundant functions. To identify genes that may be essential in the absence of EF-P in *B. subtilis*, we performed transposon mutagenesis followed by Illumina sequencing (Tn-seq). We transformed *B. subtilis* genomic DNA that had been transposon mutagenized *in vitro* (45) into wild-type and Δ*efp* cells (Fig. 1A). We pooled libraries of >450,000 colonies from each strain, then mapped and tallied the transposon insertion sites (Fig. 1B) (Dataset S1). Genes with few insertions in both wild-type and Δ*efp* cells are likely to be essential in both strains, whereas genes that have fewer insertions in Δ*efp* cells compared to wild-type are likely to be more essential in the Δ*efp* background.

**Fig 1.**
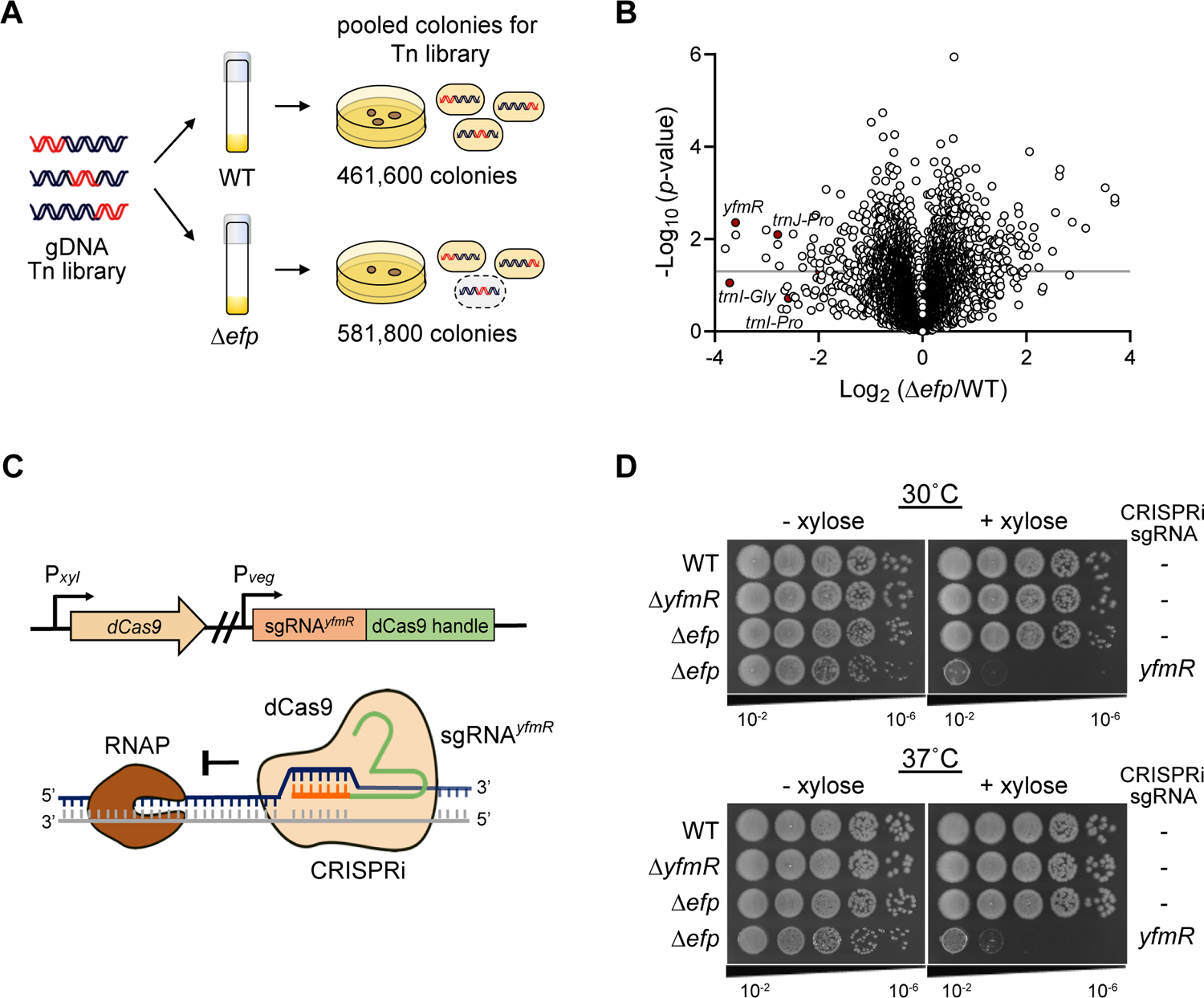
YfmR depletion is lethal to *B. subtilis* in the absence of EF-P. (A) Schematic of transposon library generation. *In vitro* transposed *B. subtilis* genomic DNA was transformed into wild-type and Δ*efp* strains. Sequencing libraries were produced in triplicate. (B) Volcano plot showing the results of Tn-seq experiment. Gray bar indicates p-value of 0.05. (C) Schematic of CRISPR interference to deplete YfmR. Guide RNA targeting *yfmR* (sgRNAyfmR) is constitutively expressed while deactivated Cas9 (dCas9) is expressed under the control of a xylose promoter. (D) *B. subtilis* Δ*efp* expressing dCas9+sgRNA*^yfmR^* fails to form colonies at 30°C and 37°C. Log-phase cultures were serially diluted and then plated with and without xylose for induction of dCas9 to deplete YfmR. Spot plates are representative of four independent experiments.

Genes with a low number of insertions in Δ*efp* cells compared to wild-type included several tRNAs that decode codons at which ribosomes stall in the Δ*efp* background (21, 46). Two of the three tRNA^Pro^ genes (*trnJ-Pro* and *trnI-Pro*) had greater than 5 times as many insertions in the wild-type background compared to the Δ*efp* background (Fig. 1B). The third tRNA^Pro^ gene (*trnB-Pro*) had very few insertions in both the wild-type and Δ*efp* background. Decoding of glycine is also strongly impacted by EF-P (21, 38, 47) and *trnI-Gly*, *trnE-Gly*, and *trnB-Gly2* averaged more than 4 times as many insertions in the wild-type than the Δ*efp* background (Fig. 1B). We did not note an apparent interaction between *efp* and other known ribosome quality control factors. *smpB*, *ssrA*, *rqcH*, *mutS2*, and *brfA* all had similarly high numbers of insertions in both the wild-type and Δ*efp* backgrounds (Dataset S1).

### Deletion of the ABCF ATPase YfmR is synthetically lethal with EF-P

One of the genes with the most significant difference in transposon insertions between the wild-type and Δ*efp* strains was *yfmR* (Fig. 1B), which encodes an ABCF ATPase (48–50). *B. subtilis* Δ*efp* cells exhibited a 10-fold reduction in insertions in *yfmR* compared to wild type, suggesting *yfmR* may be essential when *efp* is absent. *B. subtilis* encodes 4 ABCF ATPases, but only *yfmR* appeared to be essential in the Δ*efp* background. *yfmM* and *ydiF* had >100 insertions in both wild-type and Δ*efp* backgrounds, and *ykpA* had >50 insertions in both backgrounds (Dataset S1).

To test whether *yfmR* is essential in Δ*efp* cells, we first measured transformation efficiency of *ΔyfmR::kan^R^* genomic DNA into a Δ*efp* background. *ΔspoIIE::kan^R^* genomic DNA (carrying a deletion of the non-essential *spoIIE* gene) was used as a positive control. While the positive control plate had >1,000 colonies, only 2 colonies formed on the plate where *Δefp* had been transformed with *ΔyfmR::kan^R^*, and PCR revealed that both colonies maintained a wild-type copy of *yfmR* in addition to the *kan^R^* allele, indicating the *kan*^R^ allele had inserted into a different locus.

Next, we used CRISPR interference (CRISPRi) (51, 52) to deplete YfmR from Δ*efp* cells by co-expressing nuclease deficient Cas9 (dCas9) and a guide RNA targeting *yfmR (*P*_xyl_* – dCas9 P*_veg_* – sgRNA*^yfmR^*) (Fig 1C). We will refer to this strain as Δ*efp* + YfmR^depletion^. Single deletion of *yfmR* or *efp* did not significantly affect growth or colony size (Fig 1D). However, when dCas9 with sgRNA*^yfmR^* were co-expressed in Δ*efp* cells (Δ*efp* + YfmR^depletion^), these cells failed to form colonies at both 30 °C and 37 °C (Fig 1D), demonstrating that *yfmR* and *efp* are a synthetic lethal pair in *B. subtilis*.

### YfmR associates with 70S ribosomes and polysomes

To examine the association of YfmR with ribosomes, we expressed C-terminally Flag-tagged YfmR under the control of a P*_hyperspank_* promoter. We loaded cell lysate from this strain on a sucrose density gradient, then resolved an equal volume of each fraction and probed with an anti-Flag antibody. As a control, we probed these same fractions with a polyclonal antibody raised against *B. subtilis* EF-Tu (53). We detected YfmR associated with ribosomes similarly to EF-Tu, including in the polysome fractions (Fig. 2A). These data indicate that YfmR associates with actively translating ribosomes.

**Fig 2.**
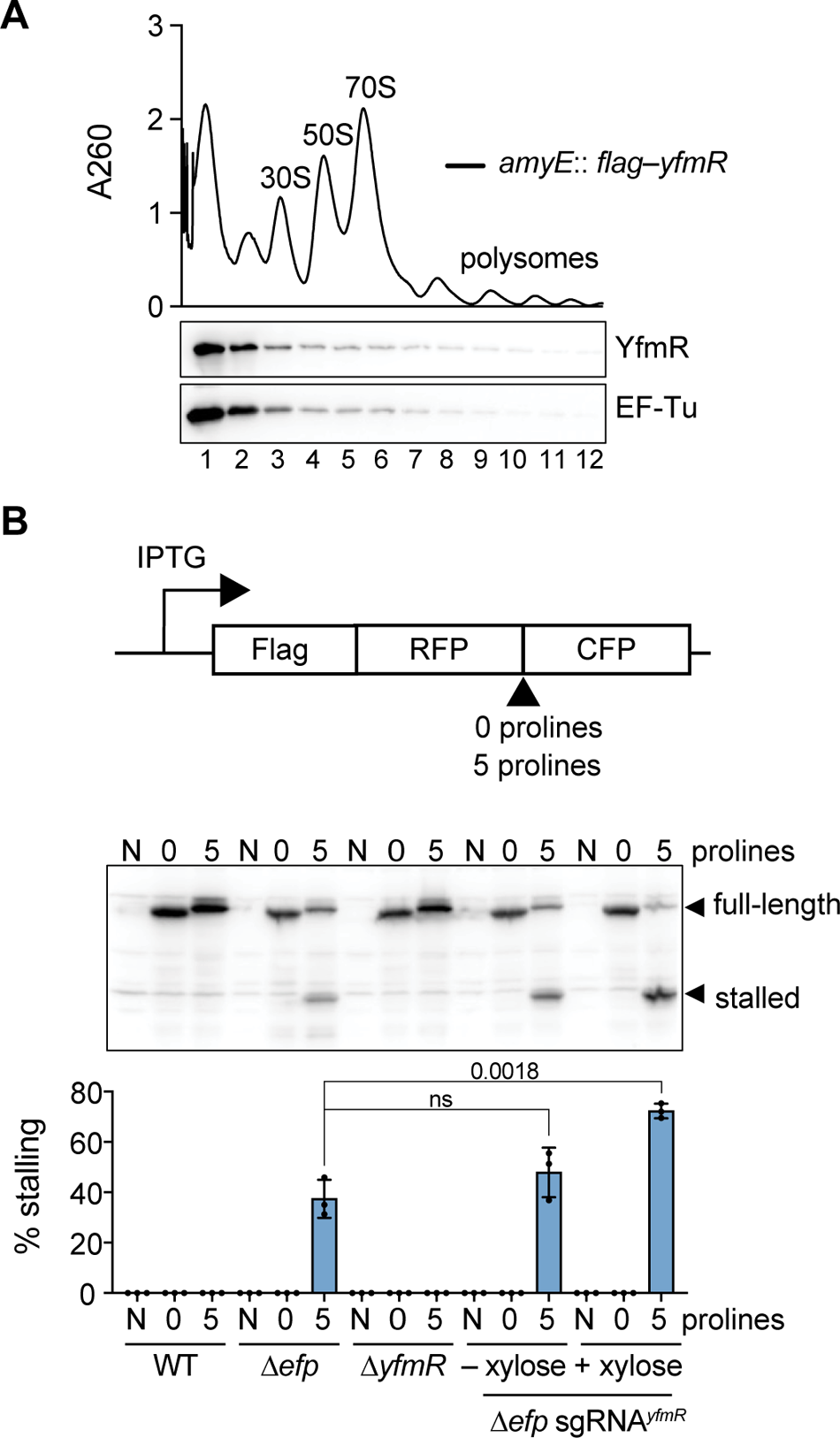
YfmR associates with ribosomes and prevents ribosome stalling at a polyproline tract. (A) Ribosome trace from sucrose density gradients and western blot of fractions probed with anti-Flag antibody to detect Flag-YfmR or anti-EF-Tu antibody to detect EF-Tu. Trace and western blots are representative of 3 independent experiments. (B) Schematic of reporter for proline stalling and western blot showing full length protein and truncated protein that results from ribosome stalling at polyproline tract. Cell lysate was harvested after 3 hours of simultaneous reporter expression and dCas9 induction to deplete YfmR. N indicates lysate from strain without reporter plasmid. 0 and 5 indicate reporter without a proline tract or with a proline tract containing 5 prolines. Error bars represent standard deviation from three independent experiments. P-value indicates the results of an unpaired two-tailed T-test.

### YfmR depletion in Δ*efp* causes severe ribosome stalling at a polyproline tract

EF-P binds to the ribosome between the E- and P-sites and assists translation of polyproline tracts and other difficult-to-translate sequences (18, 27). YfmR is closely related to Uup and EttA (50), ABCF ATPases that bind the ribosomal E-site and promote a favorable geometry for peptide bond formation (54). Therefore, we hypothesized that YfmR may play a role in rescuing ribosomes stalled at polyprolines and be functionally redundant with EF-P. To investigate this, we designed an N-terminally Flag-tagged reporter to quantify polyproline-induced stalling (Fig. 2B). An RFP-5Pro-CFP fusion was placed under the control of an IPTG-inducible promoter and integrated as a single copy into the chromosome of wild-type, *Δefp* or *Δefp* + YfmR^depletion^ backgrounds (Fig 2B). A Flag-tagged protein of the same construction but lacking the polyproline motif was used as a control. Ribosome stalling at the polyproline tract results in a peptide that is approximately 30 kDa, whereas full-length protein is approximately 58 kDa. We report ribosome stalling as a percentage of stalled peptide divided by the total of both stalled plus full-length peptide. No ribosome stalling was detected in wild-type cells whereas *Δefp* cells exhibited severe ribosome stalling at the polyproline tract (37.42 ± 7.57 % stalling). We did not detect ribosome stalling in the Δ*yfmR* single deletion (Fig. 2B). However, when YfmR was depleted from Δ*efp* cells ribosome stalling at the polyproline tract was significantly more severe than Δ*efp* deletion on its own (72.24 ± 2.9 % stalling in Δ*efp* + YfmR^depletion^)(p = 0.0018)(Fig. 2B). These data suggest that YfmR prevents ribosome stalling at a polyproline tract in the absence of EF-P.

### Depleting YfmR from *Δefp* cells reduces actively translating ribosomes and increases unengaged ribosomal subunits

To further investigate the impact of YfmR depletion in Δ*efp* cells, we used sucrose density gradient ultracentrifugation to characterize 30S and 50S ribosomal subunits, 70S ribosomes, and polysomes in wild-type, Δ*efp*, Δ*yfmR*, and Δ*efp* + YfmR^depletion^ cells. We quantified relative levels of each ribosome species by determining the area under the curve for each respective peak. Single deletions of *yfmR* or *efp* did not significantly impact levels of 30S, 50S, 70S or polysomes (Fig. 3A and Fig. S1). In contrast, when YfmR was depleted from Δ*efp* cells, 70S ribosomes and polysomes decreased by 34.2 ± 6.1 % compared to wild type (Fig. 3A)(Fig. S2)(p = 0.016). The significant decrease in 70S and polysomes indicates that there is a severe translation defect when YfmR is depleted from Δ*efp* cells.

**Fig 3.**
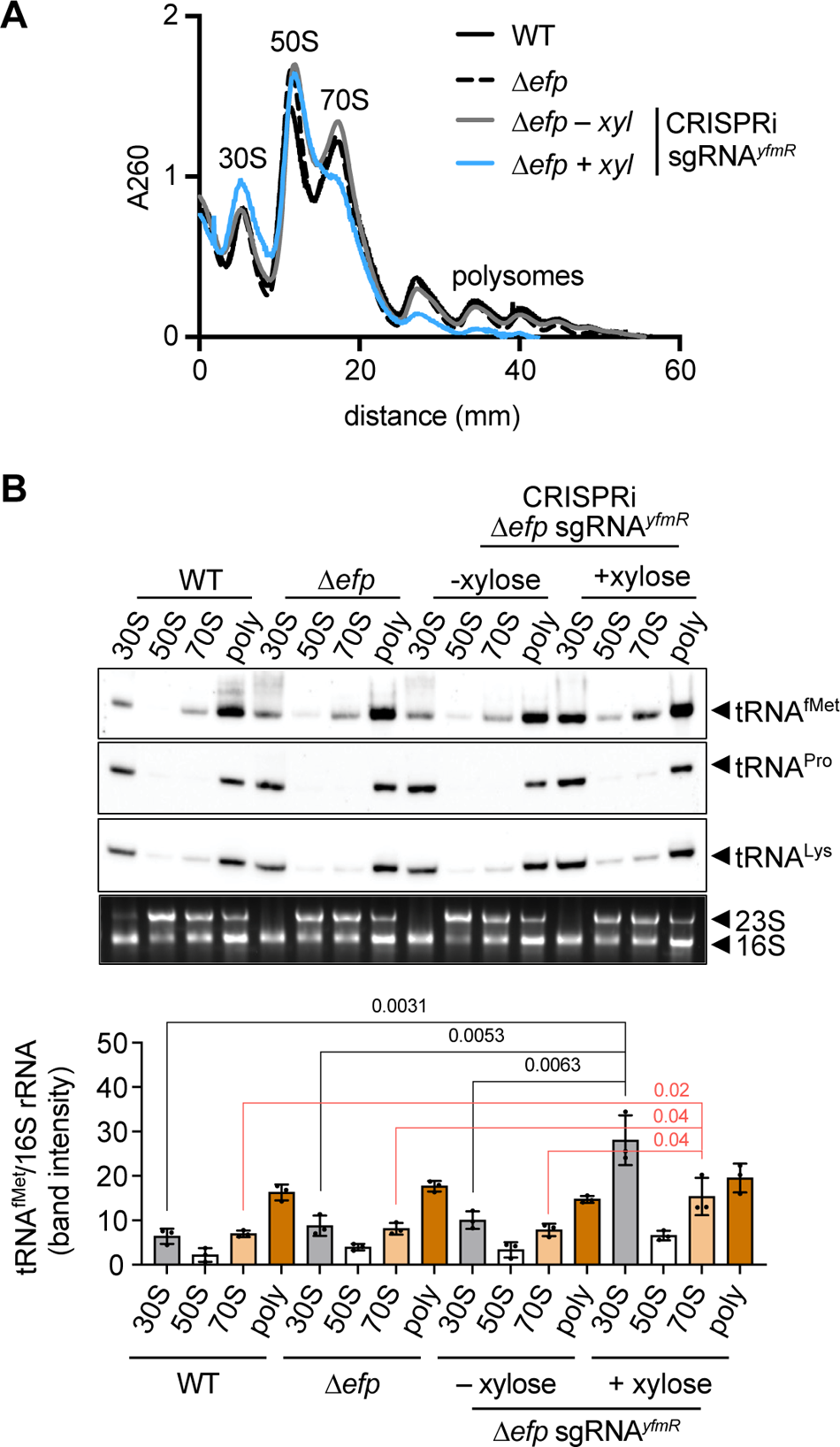
Effect of YfmR depletion on ribosome activity and tRNA association in Δ*efp* cells. Cells were grown in LB at 37°C with aeration and harvested at late-log phase. dCas9 was induced for 3 hours to deplete YfmR. (A) Sucrose density gradient ribosome traces of wild type, Δ*efp*, and Δ*efp* with and without YfmR depletion (B) Northern blot analysis of RNA precipitated from 30S, 50S, 70S, and polysome fractions using a probe specific to tRNA^fMet^, tRNA^Pro^, or tRNA^Lys^. Error bars represent standard deviation of three independent experiments. P-value indicates the results of an unpaired two-tailed T-test. See Fig S2 for quantification of tRNA^Pro^ and tRNA^Lys^.

### Initiator tRNAs are enriched in 30S and 70S ribosome fractions in *Δefp* cells when YfmR is depleted

Depleting YfmR from Δ*efp* cells resulted in decreased 70S ribosomes and polysomes and a relative increase in unengaged 30S and 50S subunits (Fig. 3A). To further explore defects in translation we assayed levels of initiator tRNA^fMet^, tRNA^Pro^, and tRNA^Lys^ associated with ribosomal subunits. We separated ribosomes by sucrose density gradient ultracentrifugation and precipitated total RNA from the 30S, 50S, 70S, and polysome peaks. The ratio of each tRNA relative to 16S rRNA was then quantified to determine levels of association with each ribosome species peak in wild-type, Δ*efp*, and Δ*efp* + YfmR^depletion^ backgrounds. Initiator tRNA^fMet^ association with the 30S and 70S peaks significantly increased when YfmR was depleted in *Δefp* (p = 0.0031 and p = 0.02) (Fig 3B). In contrast, levels of tRNA^Pro^ and tRNA^Lys^ associated with 30S or 70S did not increase in Δ*efp* cells when YfmR was depleted (Fig. 3B and Fig. S2). Increased tRNA^fMet^ association with 30S ribosomes may indicate a failure of the 50S large subunit to join, whereas increased tRNA^fMet^ association with 70S ribosomes may indicate a failure in early elongation.

### *E. coli* Uup is functionally interchangeable with YfmR

The *B. subtilis* YfmR structure predicted by Alphafold (55) is similar to *E. coli* Uup and EttA (Fig. 4A)(30, 54). Bacterial ABCFs contain two ATPase domains separated by an approximately 80 amino acid linker called the <u>P</u>-site tRNA interacting <u>m</u>otif (PtIM) (30, 56). ABCFs bind the ribosomal E-site, and their functional diversity may depend on the length and amino acid composition of the PtIM (54). EttA’s PtIM domain (residues 242-322) makes specific contacts with tRNA^fMet^ (56)(Fig. 4B) and promotes a favorable geometry for peptide bond formation during elongation (30, 54, 56, 57). YfmR has comparable overall sequence identity to both EttA (36%) and Uup (35%) but is more similar to EttA in its PtIM (Fig. 4C). YfmR and Uup both have a coiled coil C-terminal extension that is absent in EttA (50, 58).

**Fig 4.**
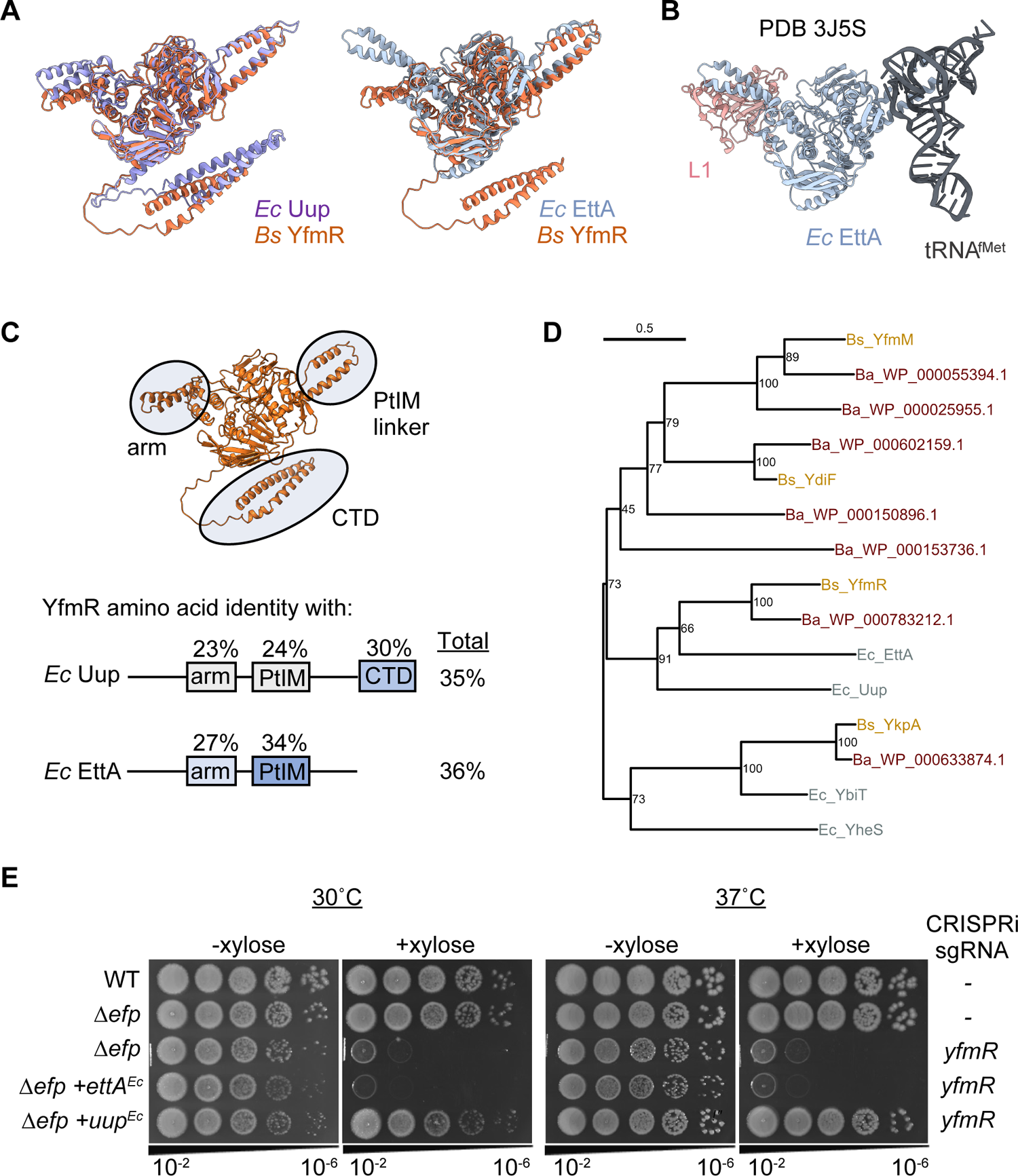
Structural, phylogenetic, and functional relationships between YfmR, Uup, and EttA. (A) Alphafold predicted structures of *B. subtilis* YfmR overlayed with *E. coli* Uup or EttA. (B) Structure of *E. coli* EttA with ribosomal protein L1 and tRNA^fMet^ (PDB 3J5S). (C) Domain homology analysis comparing YfmR to Uup and EttA. Arm domain includes YfmR residues 91 to 147. PtIM domain includes YfmR residues 239 to 318. C-terminal domain includes YfmR residues 539 to 629. (D) Phylogenetic tree of amino acid sequences of ABCFs from *B. subtilis*, *B. anthracis* and *E. coli*. Numbers at nodes indicate maximum likelihood bootstrap percentage, which indicates likelihood of a common ancestor between the two branches. (E) Spot plates with and without YfmR depletion in Δ*efp* or Δ*efp* expressing either Uup or EttA from *E. coli*.

We compared ABCFs of *B. subtilis*, *B. anthracis*, and *E. coli* to examine their likely evolutionary history. Consistent with previous reports (50), we observed that EttA and Uup share a common ancestor exclusive of the other ABCFs with a maximum likelihood bootstrap percentage of >90% (Fig 4D). YfmR from *B. subtilis* and from *B. anthracis* clustered with EttA instead of Uup, but with lower certainty (66% MLB).

To determine the functional relationship between Uup or EttA with YfmR we then tested whether Uup or EttA from *E. coli* could rescue *B. subtilis* Δ*efp* when YfmR was depleted. We expressed either Uup or EttA from an IPTG-inducible promoter integrated into the chromosome of Δ*efp* P*_veg_*–dCas9 P*_xyl_*–sgRNA*^yfmR^* (Δ*efp* + YfmR^depletion^). Expression of Uup showed a complete rescue of Δ*efp* with YfmR depletion, whereas EttA did not (Fig. 4E). These data indicate that Uup is functionally interchangeable with YfmR in *B. subtilis*.

### *B. subtilis* YfmR partially rescues *B. anthracis* Δ*efp* growth and sporulation defects

An outstanding mystery in the field is why Δ*efp* phenotypes range from modest to lethal in bacteria (38). To investigate this, we determined the phenotype of Δ*efp* in *Bacillus anthracis*, which encodes a similar number of polyproline tracts as *B. subtilis*. Whereas single deletion of *efp* in *B. subtilis* does not significantly impact growth rate and only modestly impacts sporulation efficiency (37, 44), we observed that *B. anthracis* Δ*efp* exhibits severe growth defects at both 30°C and 37°C, produces no detectable spores, and exhibits ribosome stalling at a polyproline tract (Fig. 5A, Fig. 5B, and Fig. S3). Notably, similar translation defects occur in *B. anthracis* Δ*efp* single deletion as occur in *B. subtilis* Δ*efp* when YfmR is depleted (Fig. S4A and Fig. S4B).

**Fig 5.**
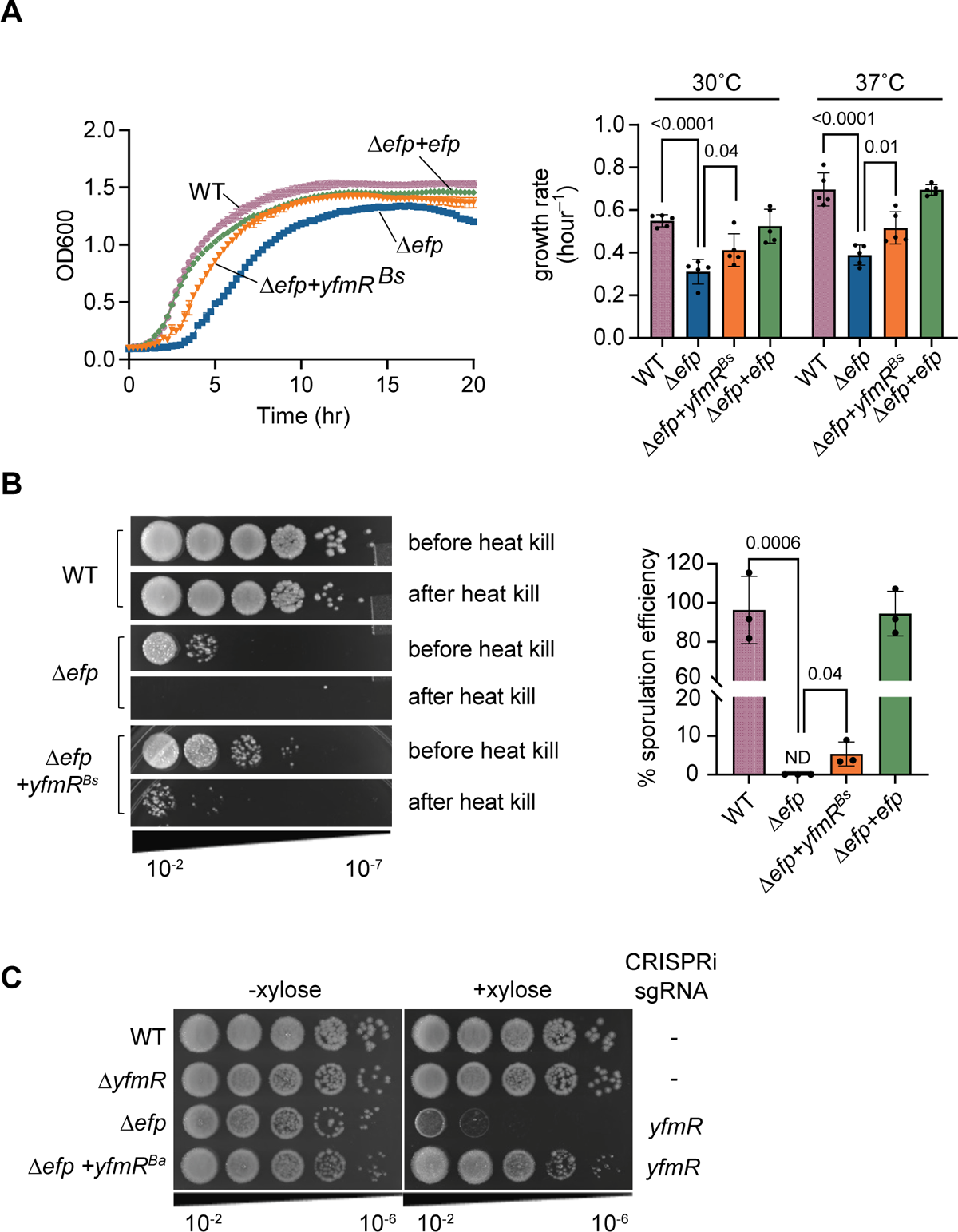
Heterologous expression of *B. subtilis* YfmR in *B. anthracis* Δ*efp* partially rescues growth and sporulation defects. A single copy of *yfmR* from *B. subtilis* (*yfmR^Bs^*) was integrated into the *B. anthracis* chromosome under the control of an IPTG-inducible promoter. (A) Growth curves of *B. anthracis* wild type, Δ*efp* and Δ*efp* expressing *B. subtilis yfmR*. Growth rates were determined at early log-phase. Error bars represent standard deviation of 5 biological replicates. P-value indicates the results of an unpaired two-tailed T-test. (B) *B. anthracis* wild type, Δ*efp* and Δ*efp* expressing *B. subtilis yfmR* was sporulated for 24 hours in DSM. Culture was serially diluted and plated before and after heating to 80°C for 20 minutes to kill non-sporulated cells. Error bars represent standard deviation of 3 biological replicates. P-values were determined by an unpaired two-tailed T-test.

To test whether *B. subtilis* YfmR could complement the *B. anthracis* growth defect we expressed IPTG-inducible YfmR from *B. subtilis* as a single copy integrated into the *B. anthracis* genome (Fig. 5A). Expression of *B. subtilis* YfmR partially but significantly rescued the growth defect of *B. anthracis* Δ*efp* at both 30°C (p = 0.04) and 37°C (p = 0.01). Expression of *B. subtilis* YfmR also partially rescued the *B. anthracis* Δ*efp* sporulation defect (Fig. 5B).

*B. anthracis* encodes a YfmR homolog (WP_000783212.1) that shares 57% amino acid sequence identity with *B. subtilis* YfmR. Structural modeling with AlphaFold (55) of *B. subtilis* and *B. anthracis* YfmR shows that they are structurally similar (Fig. S5). To determine whether *B. anthracis* YfmR is functional, we tested whether it could rescue depletion of YfmR in *B. subtilis* Δ*efp*. Expressing *B. anthracis* YfmR from an IPTG-inducible promoter in *B. subtilis* Δ*efp* + YfmR^depletion^ restored colony formation to wild type levels (Fig. 5C). These data suggest that *B. anthracis* YfmR is active and functionally redundant with *B. subtilis* YfmR. Therefore, presence or absence of an ABCF ATPase may not be sufficient to explain the variability in phenotypes of Δ*efp* mutants, and expression levels of the ABCFs must also be considered.

## Discussion

Here we identify YfmR as a translation factor that is essential in the absence of EF-P in *B. subtilis* (Fig. 1). We confirmed the essentiality of *yfmR* in Δ*efp* cells by CRISPRi (Fig. 1). YfmR associates with *B. subtilis* ribosomes, including polysomes (Fig. 2). YfmR depletion from Δ*efp* cells causes severe ribosome stalling at a polyproline tract and reduces levels of actively translating ribosomes (Fig. 2 and Fig. 3). These data support a role for YfmR in translation elongation and prevention of ribosome stalling. Additionally, we report that *B. anthracis* Δ*efp* exhibits severe growth and sporulation defects, which can be partially rescued by YfmR from *B. subtilis*, further highlighting the importance of YfmR and its functional redundancy with EF-P (Fig. 5).

Depleting YfmR from Δ*efp* cells caused more severe ribosome stalling at a polyproline tract than *efp* deletion on its own (Fig. 2B), suggesting that YfmR and EF-P have redundant functions in preventing ribosome stalling at polyprolines. Ribosome stalling at polyproline tracts in essential genes could explain the synthetic lethal phenotype of Δ*efp* Δ*yfmR*. Essential or quasi-essential genes containing a polyproline motif in *B. subtilis* include *topA*, *fmt*, and *valS*, which encode DNA topoisomerase I, methionyl-tRNA formyltransferase (FMT), and valine tRNA synthetase (ValS), respectively (59–62). FMT is of particular interest in the context of our results because it adds the N-formyl group to Met-tRNA^fMet^, which helps it to be recognized by IF2 and to resist peptidyl-tRNA hydrolysis (63, 64). The ValS tRNA synthetase is also of interest since it contains a polyproline motif that is highly conserved across eukaryotic, archaeal, and bacterial proteomes (65). It has been proposed that EF-P is universally conserved, in part, to efficiently translate this protein (65, 66).

YfmR associates with 70S ribosomes and polysomes, suggesting it plays a role in translation elongation (Fig. 2A). Reduced 70S ribosomes and polysomes, and the specific enrichment of tRNA^fMet^ associated with 30S and 70S subunits in Δ*efp* + YfmR^depletion^ further suggests that there is a failure in translation initiation or early elongation in the absence of EF-P and YfmR (Fig. 3). EF-P can promote early elongation at some amino acids *in vitro* (26, 28). Therefore, further experiments are required to determine whether YfmR can also facilitate early elongation or whether the decreased translation we observe is an indirect result of cell death or the failure to produce an essential translation factor containing a polyproline motif.

YfmR is closely related to both Uup and EttA (Fig. 4) (50). Heterologous expression of *E. coli* Uup restored growth of *B. subtilis* Δ*efp* cells when YfmR was depleted (Fig. 4E), suggesting that YfmR and Uup have a similar function *in vivo*. Uup does not stimulate peptide bond formation in a tripeptide assay using fMet-Phe-Lys (57). However, it is possible that, like EF-P, Uup does not stimulate peptide bond formation between fMet-Phe, but may stimulate peptide bond formation between fMet and other amino acids (26, 29). Indeed, a recent preprint showed that Uup increased production of a peptide encoding a polyproline tract *in vitro* (67). Moreover, Uup-EQ2 has similar binding affinity for both initiating and elongating ribosomes (54), further suggesting that Uup may function in elongation. Uup also likely plays a role in ribosome biogenesis since Δ*bipA* Δ*uup* cells accumulate immature 50S subunits (50, 68). Since EF-P does not play a role in ribosome biogenesis, our data suggest that the essential function of YfmR/Uup in the absence of EF-P is not ribosome biogenesis, but prevention of ribosome stalling.

ABCFs are near-universally conserved in bacterial genomes, with an average of four paralogs encoded per genome (50). The diverse functions of these highly conserved proteins are just beginning to be determined. Many important questions remain regarding their evolution, functional overlap, regulation, and mechanisms of action. Our work reveals the physiological importance of one of the most highly conserved ABCFs (YfmR/Uup) in maintaining cell viability and preventing ribosome stalling *in vivo*.

## Data Availability Statement

Sequencing data are available at Gene Expression Omnibus accession GSE249203. An Excel table containing the results of the Tn-seq can be found in supplemental information (Dataset S1). Strains and plasmids are available upon request.

## Methods

See SI Appendix, Methods for detailed descriptions. Strains, plasmids, and primers are found in Table S1, S2, and S3.

### Strains and media

All strains were derived from *B. subtilis* 168 *trpC2* and *B. anthracis* Sterne 34F2. *B. subtilis* and *B. anthracis* were grown in LB media at 30°C and 37°C as indicated. Sporulation was induced in Difco Sporulation Media (DSM) with aeration for 24 hours at 37°C.

### Strain Construction

Markerless *efp* deletion in *B. anthracis* was made by two-step recombination as previously described (69). Gene deletions in *B. subtilis* were made by transforming genomic DNA from a BKK strain deletion in the lab’s wild-type *B. subtilis 168 trpC2* (HAF1) (59). Δ*efp* + YfmR^depletion^ was constructed by expressing dCas9 under the control of a xylose-inducible promoter (51) while expressing a guide RNA targeting *yfmR* from a vegetative promoter in the Δ*efp* background.

### Tn-seq library and sequencing

Magellan6x *in vitro* transposed gDNA prepared from wild-type *B. subtilis* 168 (45) was transformed into wild-type or Δ*efp B. subtilis*. Transposon insertions were sequenced on the Illumina MiSeq platform (v3 150 bp kit) at the Cornell University Biotechnology Resource Center.

### Measuring proline stalling

P*_hyperspank_*–3xFLAG–RFP–linker–CFP was integrated at *thrC* in the wild-type, *Δefp* or *Δefp* + YfmR^depletion^ background. The linker contained 0 or 5 prolines. The proline staller was induced with 1 mM IPTG at the same time the sgRNA*^yfmR^* was induced with 5% xylose. Induction of the reporter and depletion of YfmR proceeded for 3 hours before harvesting cells.

### Sucrose gradient fractionation

Wild-type, *Δefp* or *Δefp* + YfmR^depletion^ cells were grown in LB starting from an OD of 0.05. Xylose was added to 5% to induce sgRNA*^yfmR^* and cells were harvested 3 hours later. Clarified cell lysates were normalized to 1500 ng/µl and loaded onto 10 – 40% sucrose gradients.

### Northern blotting and RNA gels

RNA from sucrose gradient fractions were precipitated and normalized to 1 µg RNA. RNA was resolved on a TBE-Urea gel and transferred to a positively-charged nylon membrane. RNA was probed with a biotinylated oligo specific for tRNA^fMet^, tRNA^Pro^, or tRNA^Lys^. Oligo sequences are in Table S3.

### Structure modeling and sequence comparison of ABCFs

The *E. coli* EttA structure was obtained from Protein Data Bank (PDB 3J5S) (30). YfmR structures for *B. subtilis* and *B. anthracis* were generated with AlphaFold (55). Molecular graphics and analysis were performed with ChimeraX v1.5 (70). ABCF protein sequences were downloaded from NCBI *E. coli* str. K-12 substr. MG1655 (NC_000913.3) and *B. subtilis* subsp. subtilis str. 168 (NC_000964.3) reference genomes as annotated by the NCBI Prokaryotic Genome Annotation Pipeline (71).

## Acknowledgements

We are grateful to lab members Katrina Callan and Kevin England for feedback and discussion of the work. We are grateful to Scott Stibitz and Roger Plaut for sharing pRP1099, pRP1028, pSS1827 and for advice on constructing gene deletions in *B. anthracis* Sterne. We are grateful to Christopher Johnson and Alan Grossman for sharing their transposon library. HAF, HRH, LW, DDT and CRP were supported by NIH grant R35GM147049. LW was supported by an undergraduate research fellowship from the Cornell Institute of Host-Microbe Interactions and Disease.

## SI Materials and Methods

### Strains and media

All strains were derived from *B. subtilis* 168 trpC2 and *B. anthracis* Sterne 34F2 and are listed in Table 1. *B. subtilis* and *B. anthracis* were grown in LB media (10 g Tryptone, 5 g yeast extract, and 5 g NaCl per liter) at 30°C and 37°C as indicated. *B. anthracis* strains were cultured in Brain Heart Infusion (BHI) for conjugation. *E. coli* strains were cultured in LB. Antibiotics were used at final concentrations of 100 µg/mL ampicillin (58), and 10 µg/mL kanamycin, 100 µg/mL spectinomycin, and 1x MLS (1 µg/mL erythromycin and 25 µg/mL lincomycin).

### Construction of *B. anthracis* Δ*efp*

Primers are listed in Table S2. A DNA fragment (Twist Biosciences) containing 700-bp regions flanking the *efp* open reading frame along with 30 bp of homology to pRP1028 at the 5’ and 3’ ends was cloned into pRP1028 (1) at the BamHI restriction site by Gibson assembly (2) to create plasmid pHRH28. pHRH28 was conjugated into *B. anthracis* Sterne by tri-parental mating as described previously (1). Briefly, the strain harboring the donor plasmid was mixed with helper plasmid pSS1827 (3) and recipient *B. anthracis* Sterne by mixing on BHI plates and incubating at room temperature overnight. The mixture was scraped and spread onto BHI agar media supplemented with 250 µg/mL spectinomycin and 60 units of Polymyxin B. Integrants were restreaked on BHI plates containing spectinomycin. A second tri-parental mating was then performed to transform the integrants with plasmid pRP1099 harboring the SceI nuclease. Donor, helper, and recipient strains were mixed and the mating proceeded at 37°C for 8 hours. Since the previously integrated donor plasmid pRP1028 contains an SceI endonuclease target site, introducing the new donor plasmid pRP1099 carrying the SceI endonuclease facilitates *B. anthracis* chromosome recombination. Spectinomycin sensitive colonies were screened for *efp* deletion by PCR using primers HRH26 and HRH27. A colony containing the deletion was cured of pRP1099 to yield Δ*efp B. anthracis*. The deletion was further confirmed by whole genome sequencing (MiGS SeqCenter).

### Complementation of *efp* and *yfmR* in *B. anthracis*

We first constructed a plasmid to integrate genes under the control of an IPTG inducible promoter as a single copy at the *sacA* locus in *B. anthracis*. A DNA fragment (Twist Biosciences) containing 650-bp regions up- and downstream of *sacA* and with 30 bp of homology to pRP1028 at 5’ and 3’ end was cloned into pRP1028 by Gibson assembly, producing pHRH190. The DNA fragment was designed with a BamHI site in between the two fragments. *B. anthracis efp* was cloned into pDR111 by assembling pDR111 digested with HindIII and SphI and a Twist fragment containing the *efp* coding sequence to make pHRH101. The region between BamHI and EcoRI of pDR111 containing the *hyperspank* promoter, *efp*, and *lacI* were then amplified using primers HRH54 and HRH67 and cloned into pHRH190 by Gibson assembly. *B. subtilis yfmR* was cloned into pDR111 with primers (HRH157 and HRH158) and then the region between BamHI and EcoRI was moved into pHRH190 using the same method as for the *efp* complementation. Triparental mating was used to deliver the resulting plasmids into *B. anthracis* Δ*efp*. The resulting transconjugant was mated with pSS1827 and DH5α pRP1099 for homologous recombination to create a complementation strain that was Δ*efp* + *efp* (HRH318). Insertion of *efp* at *sacA* for complementation was PCR-confirmed with primers HRH60 and HRH61. The plasmid pRP1099 harboring SceI endonuclease was cured from the complemented Δ*efp B. anthracis*.

### Complementation of Δ*efp* + YfmR depletion

*yfmR* from *B. anthracis* was amplified with primers HRH197 and HRH198. *E. coli* MG1655 *ettA* was amplified with primers HRH199 and HRH200. *E. coli uup* was amplified with primers HRH201 and HRH202. The fragments were Gibson assembled into pDR111 digested with HindIII and SpHI to create pHRH925, pHRH983, and pHRH987. The resulting plasmids were amplified with primers HRH193 and HRH194 to amplify the region containing the P*_hyperspank_* promoter and moved to pJMP3 (4) digested with EcoRI. These resulting plasmids were linearized with ScaI and transformed into *B. subtilis* for double crossover at *thrC*.

### Sporulation

Sporulation was induced in Difco Sporulation Media (DSM) with aeration for 24 hours at 37°C. Cultures were serially diluted in Tbase with 1 mM MgSO_4_ and 10 µL was spot plated on LB agar plates before and after heating at 80°C for 30 min to kill non-sporulated, vegetative cells. Sporulation efficiency was calculated as the percentage of colony forming units before and after heating.

### Tn-seq library and sequencing

Magellan 6x *in vitro* transposed gDNA prepared from wild-type *B. subtilis* 168 (5) was transformed into wild type or Δ*efp B. subtilis*. 40 µg of gDNA was transformed into 10 mL of cells made naturally competent (6). Over 400,000 colonies from each library were pooled in 200 mL of LB. gDNA was extracted and sequencing libraries prepared in triplicate from 5 mL each of culture as described previously (39). Briefly, 6 µg of gDNA was digested with 3 units of MmeI for 2.5 hours at 37°C, then treated with 6 units of antarctic phosphatase for 1 hour at 37°C. DNA was then extracted with 300 µl phenol:chloroform:isoamyl alcohol (25:24:1) and precipitated with 50 µL 3M sodium acetate and 500 µL isopropyl alcohol. Dried pellets were dissolved in 27.5 µL 2 mM Tris-HCl pH 8.5. The DNA was ligated with one volume of 200 µM annealed adapters HRH-Tn1 and HRH-Tn2 in 1 mM Tris-Cl pH 8.5. The ligated DNA was purified with the Qiagen PCR purification kit and amplified with the primers HRH-Tn3 and HRH-Tn4. The resulting 136 base pair band was gel extracted and sequenced on the Illumina MiSeq platform (v3 150 bp kit) at the Cornell University Biotechnology Resource Center. Reads were mapped to the *B. subtilis* 168 trpC2 genome (NCBI, NC_000964.3) and tallied using Geneius Prime (Biomatters) (Dataset S1). A volcano plot was created with Graphpad Prism using a two-tailed T-test to determine P-value.

### CRISPRi knockdown

To construct an sgRNA targeting YfmR, pJMP2 (4) was amplified with primer HRHsgRNA-YfmR containing sgRNA sequence 5’-GTTGATTTTCCTGTTCCGT-3’ targeting yfmR and the reverse primer HRH175. The amplified construct was treated with DpnI for 1 hour at 37°C to remove the template plasmid. The PCR product was gel extracted and reassembled by Gibson assembly to create pJMP2-sgRNAyfmR. The reaction was transformed into DH5α and the sequence was confirmed with Sanger sequencing. pJMP1 carrying CRISPRi dCas9 under the control of a xylose-inducible promoter was transformed into wild-type *B. subtilis* and *Δefp* for integration at *lacA* (4). pJMP2-P*_veg_* – sgRNAyfmR was transformed into the resulting strains for integration at *amyE*. The resulting *B. subtillis* CRISPRi knockdown strains were grown overnight in the absence of xylose and then back diluted to an OD600 of 0.05 in LB. The culture was serially diluted and 10 µL of each dilution was spotted onto LB agar or onto LB agar + 5% xylose.

### Proline stalling reporter

For *B. anthracis* the reporter plasmid was first generated by liberating the SceI endonuclease encoding gene with EcoRI and SalI from pRP1099 (1) and subsequently removing CFP with inverse PCR using primers HRH73 and HRH74. Next, the fragment P*_hyperspank_*–MCS–lacI of pDR111 was amplified with HRH36 and HRH37 and cloned into the plasmid by Gibson assembly, producing pHRH270. The plasmid was cut with HindIII and SphI, and the fragment containing P*_hyperspank_*–3xFLAG–RFP–linker–CFP was ordered as a Twist Bioscience gene fragment and cloned in by Giboson assembly to produce pHRH551. The linker region was modified to have either three-polyprolines 5’-CCA CCT CCG-3’ (pHRH609), or five polyproline motifs 5’-CCA CCA CCA CCA CCC-3’ (pHRH605). Modified linker regions were purchased as a gene fragment from Twist Biosciences. The resulting reporter plasmids were sequenced with Plasmidsaurus and conjugated into *B. anthracis* strains by triparental mating as described above and selected on kanamycin. *For B. subtilis*, the fragment P*_hyperspank_*–3xFLAG– RFP–linker–CFP was amplified from pHRH609 (for the 3 proline motif), or pHRH605 (for the 5 proline motif) with primers HRH36 and HRH37 and cloned into pDR111 by Gibson assembly.

Next, we amplified the region containing P*_hyperspank_*–3xFLAG–RFP–linker–CFP with primers HRH193 and HRH194 and Gibson assembled into pJMP3 cut with EcoRI. These reporter plasmids were linearized with ScaI and transformed into *B. subtilis* for double crossover at *thrC*.

### Western blotting

Overnight cultures of strains carrying the stalling reporter were normalized to an OD600 of 0.05 and induced with 1 mM IPTG. The cultures were grown up to OD600 0.6 and 1.2 at 37°C to measure stalling at mid- and late-log phase. 0.6 OD units of culture were pelleted and resuspended in lysis buffer (10 mM Tris pH 8, 50 mM EDTA, 1 mg/mL lysozyme). After lysis at 37°C for 10 minutes, SDS loading dye was added and lysate heated to 90°C for 5 minutes. Proteins were resolved on a 12% SDS-PAGE gel run at 150 V for 70 minutes. Proteins were transferred to PVDF membrane (BioRad) at 300 mAmps for 100 minutes. The membrane was blocked with 3% bovine serum albumin (BSA) for 1 hour at room temperature. 1 µL of anti-FLAG antibody conjugated to Horseradish Peroxidase (Sigma SAB4200119) in 10 mL of PBS-T was incubated overnight at 4°C. The blot was washed 3 times in PBS-T and developed by using ECL substrate and enhancer (Biorad 170-5060) for detection of full length and truncated bands indicating ribosome stalling. For western blotting against HPF, 10 µL of each fraction collected from a sucrose density gradient was combined with 4X SDS-loading buffer and separated by SDS-PAGE. Protein was transferred to PVDF membrane at 300 mAmps for 90 minutes, blocked with 3% BSA and incubated with a polyclonal antibody raised against *B. subtilis* HPF (7) for 1 hour at room temperature. The blot was then washed 3 times in PBS-T and incubated with secondary anti-rabbit antibody conjugated to HRP for 30 minutes at room temperature. The blot was then washed 3 times in PBS-T and developed with ECL (BioRad 170-5060).

### YfmR ribosome association

C-terminally Flag-tagged *yfmR* was cloned into pDR111 under the control of the *hyperspank* promoter. Cells were grown to late log-phase, lysed, and loaded on a 10%-40% sucrose gradient in gradient buffer (20 mM Tris-acetate [pH 7.44°C], 60 mM ammonium chloride, 7.5 mM magnesium acetate, 6 mM β-mercaptoethanol, 0.5 mM EDTA) and centrifuged for 3 hours at 4°C at 30,000 RPM in an SW-41 rotor. Equal volumes of gradient fractions were separated by SDS-PAGE and probed with an anti-Flag antibody (Sigma A8592) or with polyclonal antibody raised against *B. subtilis* EF-Tu (8).

### Sucrose gradient fractionation

*B. subtilis* and *B. anthracis* cells were grown overnight at 37°C and inoculated to an OD600 of 0.05 in 50 ml LB. The LB media was supplemented with 1 mM IPTG for the EF-P complementation strain. Log phase culture (∼OD 0.6) was collected by centrifugation at 8000 rpm for 10 minutes (Beckman Coulter Avanti J-15R, rotor JA-10.100). The pellets were resuspended in 200 µl gradient buffer containing 20 mM Tris (pH 7.4 at 4°C), 0.5 mM EDTA, 60 mM NH_4_Cl, 7.5 mM MgCl_2_, and 6 mM BME. The cell resuspension was lysed using a homogenizer (Beadbug6, Benchmark) by four 30 s pulses at speed 4350 rpm with chilling on ice for 3 min between the cycles and clarified by centrifugation at 21,300 rcf for 20 min (Eppendorf 5425R, rotor FA-24×2). Clarified cell lysates were normalized to 1500 ng/µl and loaded onto 10 – 40% sucrose gradients in gradient buffer. Ultracentrifugation separated ribosome species at 30,000 rpm for 3 hours at 4°C (Beckman Coulter, rotor SW-41Ti). Gradients were collected using a Biocomp Gradient Station (BioComp Instruments) with A260 continuous readout (Triax full spectrum flow cell). The area under each peak for 30S, 50S, and 70S and the several peaks for polysomes, was quantified using Prism GraphPad.

### Northern blotting and RNA gels

RNA from 0.5 mL of sucrose gradient fractions were precipitated with 1.5 mL of 200 proof ethanol and incubated overnight at −20°C. The sample pellets were collected by centrifugation at 14,500 rpm for 30 minutes at 4°C. After drying the pellets for 20 minutes at 4°C, pellets were dissolved in 80 μL of nuclease-free water and normalized to 1 µg RNA in 5 µL nuclease-free water, then mixed with 5 µl 2x RNA loading dye (NEB). To normalize loading for northern blots by rRNA, 1 µg RNA for each sample was heated at 65 °C for 5 minutes and loaded onto 1.5% TBE agarose supplemented with 3 µL of 10 mg/ml ethidium bromide at 90 V for 1 hour. Bands were quantified and normalized for total rRNA. For detecting each tRNA species, RNA normalized as described for each sample was heated at 90°C for 4 minutes and loaded onto 10% TBE urea gel (Biorad), then run at 100 V for 110 minutes. The sample RNAs were transferred to Brightstart plus nylon membrane (Invitrogen) at 20 V for 16 hours at 4°C. The membrane was crosslinked with UV Stratalinker 1800 at 1200 × 100 µJ/cm2.

The crosslinked membrane was pre-hybridized with 20 mL hybridization buffer (Invitrogen) at 45°C for 1 hour. 10 µL of 20 µM 5’ biotinylated probe was added to the pre-hybridized membrane, then incubated at 45 °C for 17 hours. The membrane was washed twice with a buffer containing 1x PBS and 0.5% SDS at 45 °C, shaking at 60 rpm. Next, the blot was washed with a buffer containing 2x sodium saline citrate (SSC, VWR) and 0.5% SDS at 45°C, shaking at 60 rpm. The washed membrane was blocked with 20 mL of 3% BSA in 1x PBS + 0.5% SDS at room temperature for 1 hour. We then added 2 µL 1.25 mg/ml streptavidin-HRP (Thermo Scientific) to the 20 mL blocking solution and incubated for 1 hour. The membrane was washed with wash buffer (1x PBS and 0.5% SDS) three times and the blot was developed with 0.8 mL ECL reagent (BioRad) and imaged using a ChemiDoc MP imaging system (BioRad). tRNA sequences for probe design were obtained from GtRNAdb version 2.0 (9). tRNA^Lys^ sequence was published previously (10). Band intensity for the respective tRNA was compared to band intensity of the 16S rRNA in the same sample to determine the ratio of tRNA relative to the amount of small ribosomal subunit.

### Structure modeling and sequence comparison of ABCFs

The *E. coli* EttA structure was obtained from Protein Data Bank (PDB 3J5S) (11). YfmR structures for *B. subtilis* and *B. anthracis* were generated with AlphaFold (12). Molecular graphics and analysis were performed with ChimeraX v1.5 (13). ABCF protein sequences were downloaded from NCBI *E. coli* str. K-12 substr. MG1655 (NC_000913.3) and *B. subtilis subsp. subtilis str.* 168 (NC_000964.3) reference genomes as annotated by the NCBI Prokaryotic Genome Annotation Pipeline (14). Homologs in *B. anthracis* str. Sterne (NZ_CP009541.1) were identified by searching *B. subtilis* ABCF protein sequences (YfmM, YdiF, YfmR, YkpA) against all annotated *B. anthracis* protein sequences using HMMER v3.3 (phmmer) (15). Hits with E-values below 1e-50 and scores above 175 were used for further analyses. ABCF protein sequences were aligned using MAFFT v7.453 (16). The alignment was used to build a maximum likelihood phylogenetic tree in RAxML v8.2.12 using the PROTGAMMAAUTO and autoMRE options (17). The tree was midpoint rooted using the phangorn package v2.11.1 (18) and visualized using ggtree v3.6.2 (19).

---

**Fig. S1.**
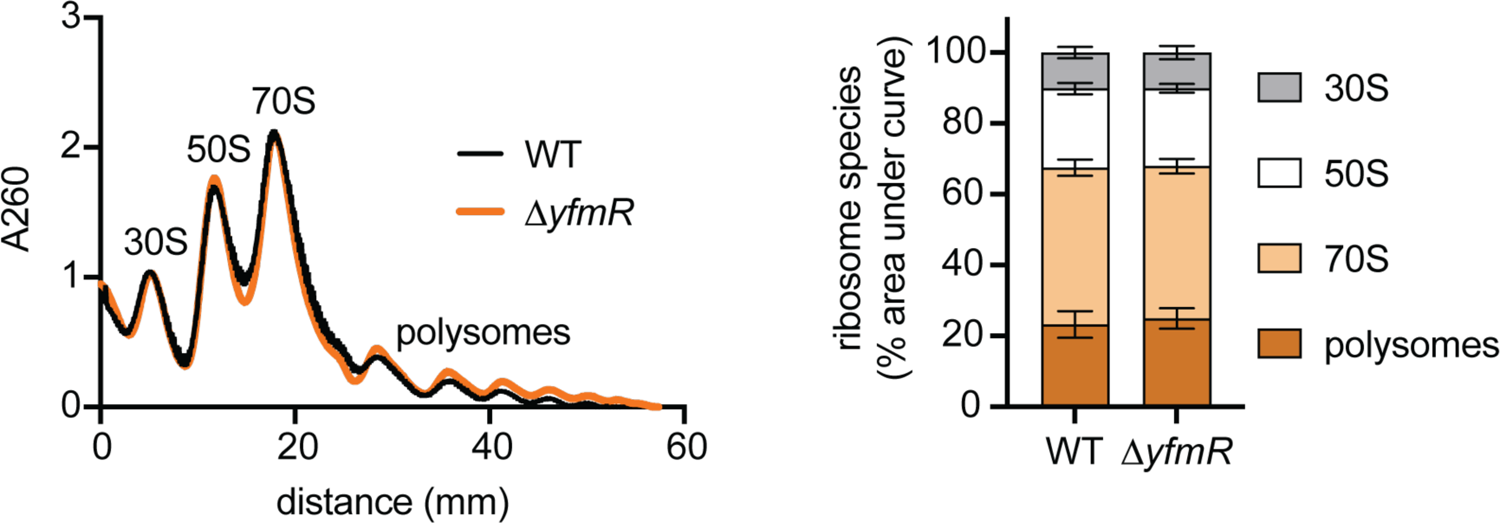
Deleting *yfmR* does not affect ribosome biogenesis. *B. subtilis* wild type and Δ*yfmR* were grown to late-log phase. Cleared lysate was loaded on a 10-40% sucrose gradient. (Left) Sucrose density gradient traces of cell lysate from wild-type and Δ*yfmR B. subtilis*. (Right) Quantification of 30S, 50S, 70S, and polysomes as determined by area under the curve. Error bars represent standard deviation of three biological replicates. No significant difference was observed between wild type and *ΔyfmR*.

**Fig. S2.**
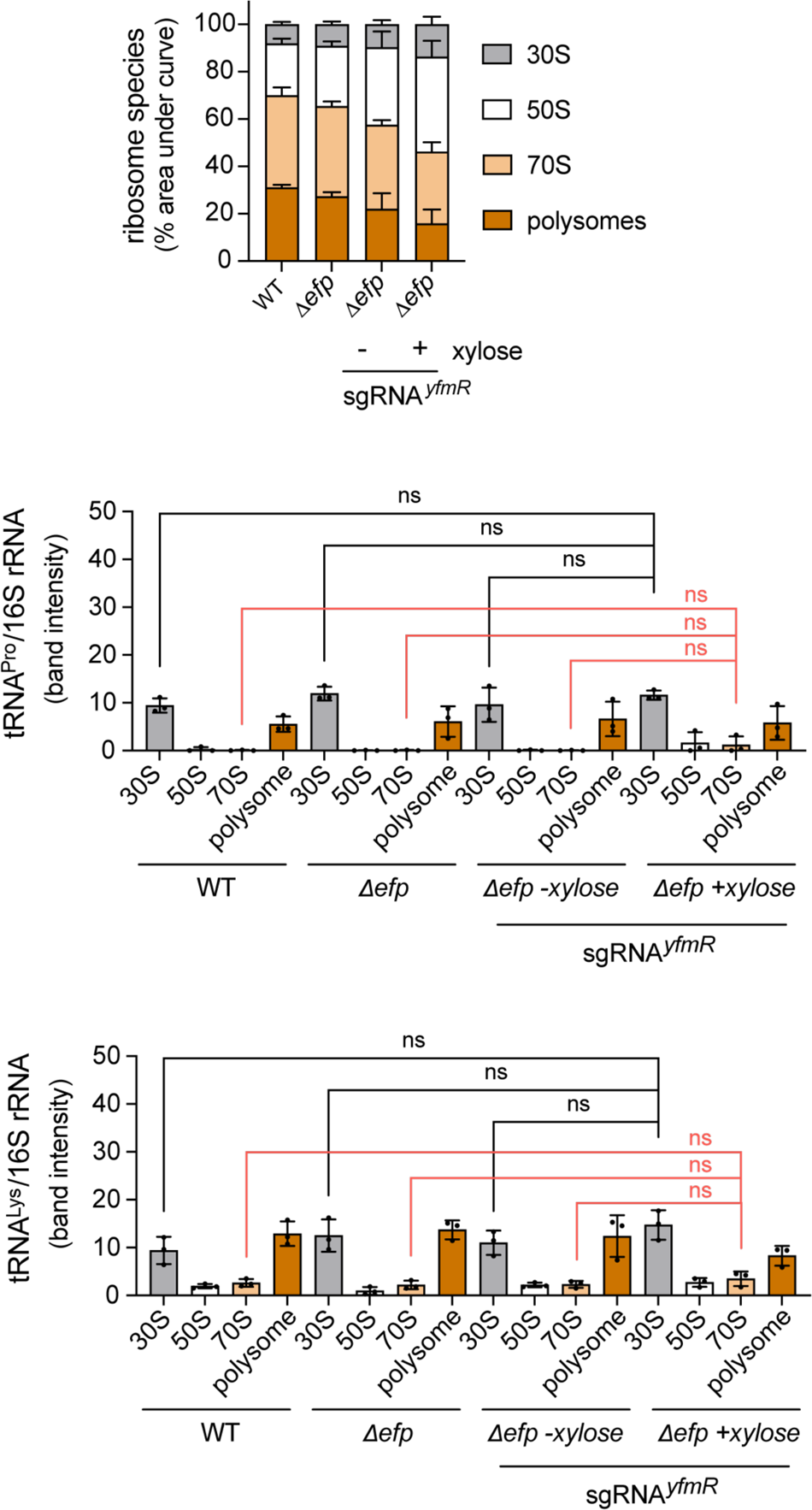
Quantification of the effect of YfmR depletion on ribosome composition and tRNA association. Cells were harvested at late log phase. dCas9 was induced for 3 hours to deplete YfmR. (A) Quantification of area under the curve for sucrose density gradients of wild-type, Δ*efp*, and Δ*efp* cells with and without YfmR depletion. (B) Quantification of tRNA^Pro^ and tRNA^Lys^ association with various ribosomal subunits as determined by northern blot. Error bars represent standard deviation of three independent experiments. P-value indicates the results of an unpaired two-tailed T-test.

**Fig. S3.**
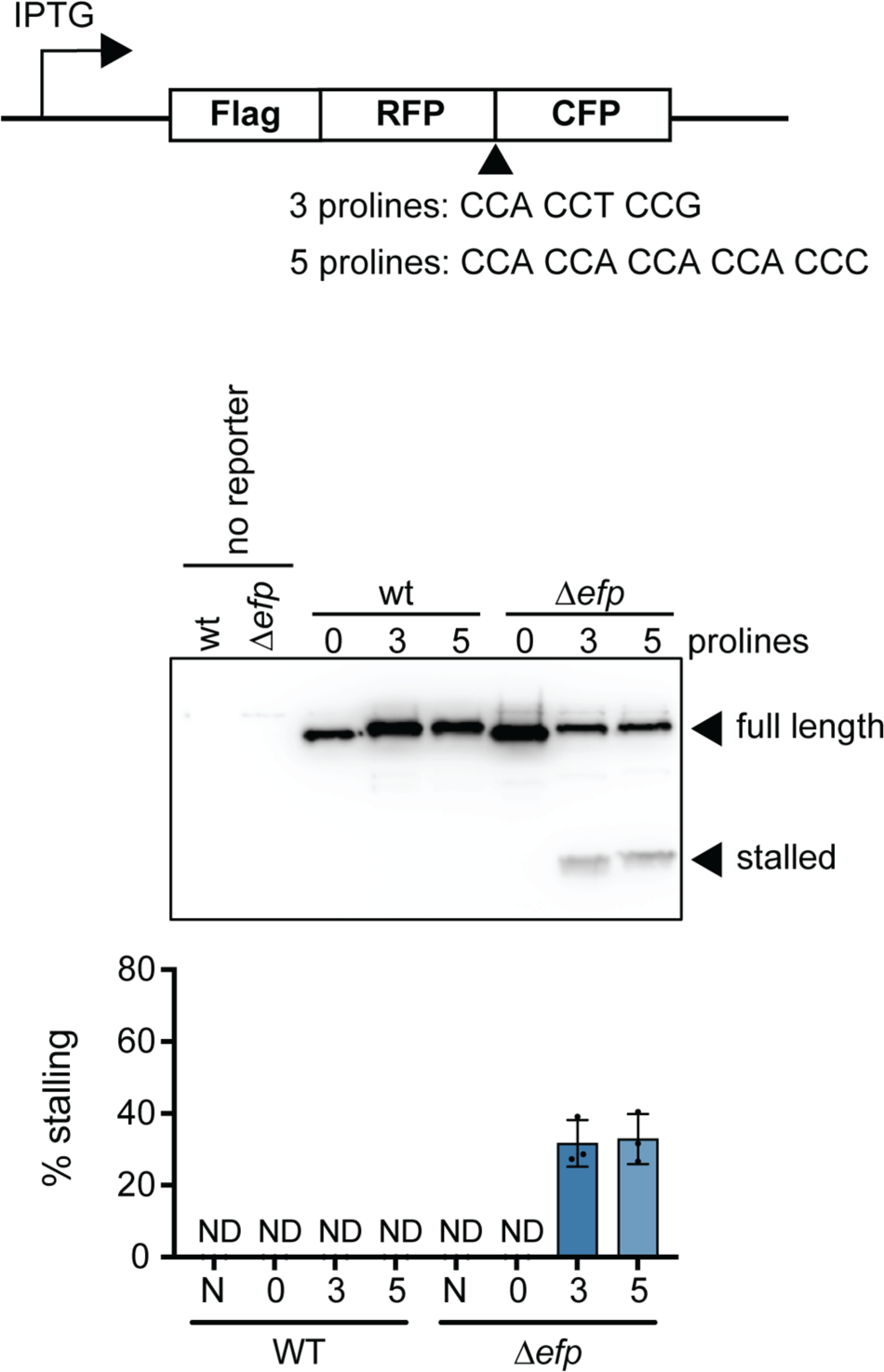
Measurement of ribosome stalling on polyproline motifs in *B. anthracis* wild type and Δ*efp*. A stalling reporter was constructed with three (CCA CCT CCG) or five (CCA CCA CCA CCA CCC) consecutive prolines in between an RFP and CFP fusion. The 3-proline stalling sequence is the motif from ValS (valyl-tRNA synthetase) of *B. anthracis*. Ribosome stalling at each motif was measured by western blot using anti-Flag antibody. Percent stalling was determined by measuring the ratio of stalled band intensity to total band intensity for both full-length and stalled peptide. Stalling levels were similar for both 3x prolines (32.9±7%) and 5x prolines (31.7±6.5%). ND indicates not detectable. Error bars represent standard deviation of 3 biological replicates.

**Fig. S4.**
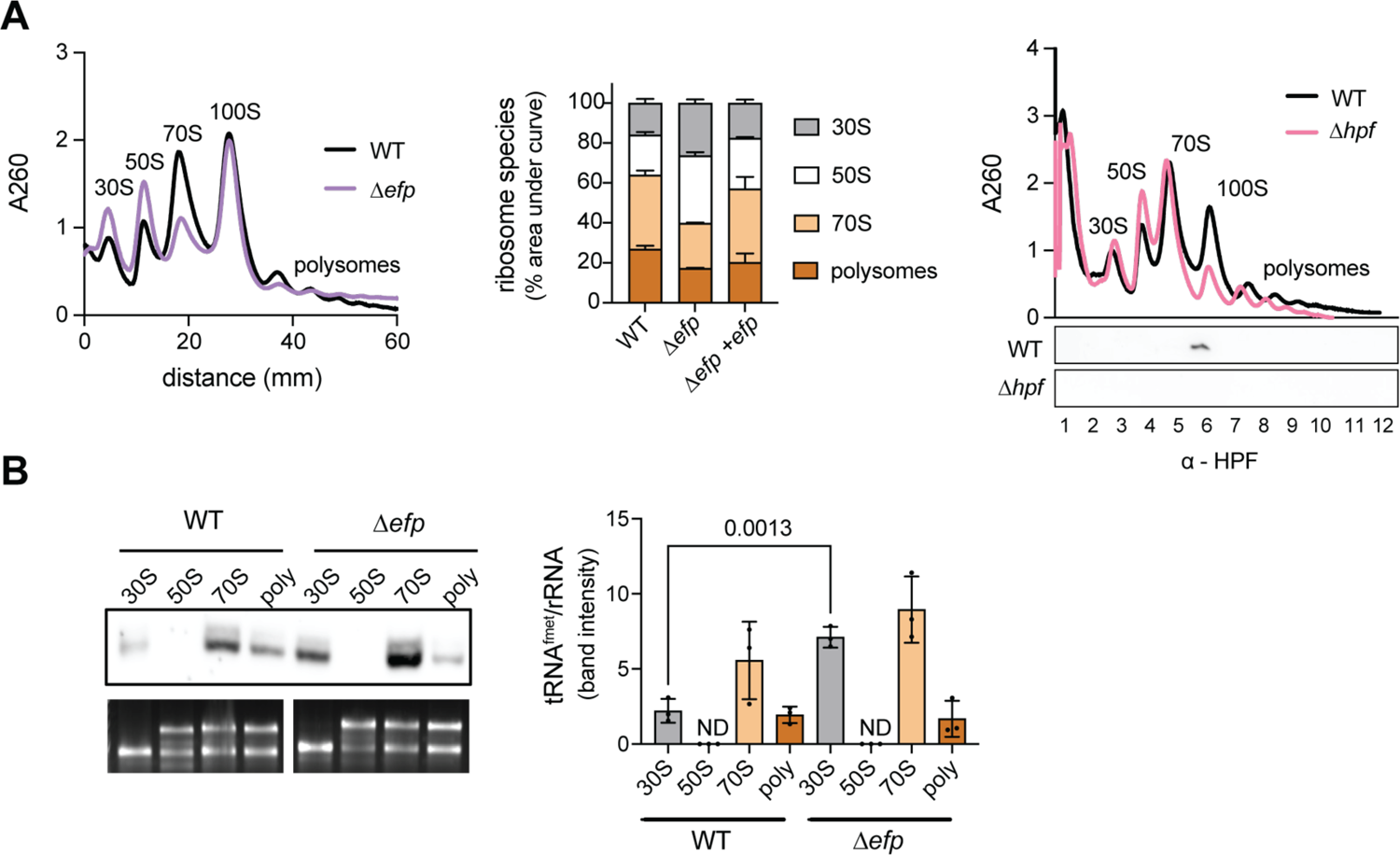
Effect of *efp* deletion on *B. anthracis* ribosomes. (A) Wild type and Δ*efp B. anthracis* were grown to late log phase at 37°C in LB media. Ribosome gradients from sucrose gradients are shown with quantification of area under the curve for each peak. Error bars represent standard deviation of 3 biological replicates. At right, sucrose gradient and western blot confirming that the 100S peak contains HPF-bound ribosome dimers. Wild type and Δ*hpf B. anthracis* were grown to late log-phase at 37°C in LB media. Gradient fractions were probed with antibody raised against *B. subtilis* HPF. (B) Northern blot to assay levels of tRNA^fMet^ associated with ribosomal subunits and polysomes. Error bars represent standard deviation of 3 biological replicates. P-values indicate the results of an unpaired two-tailed T-test.

**Fig. S5.**
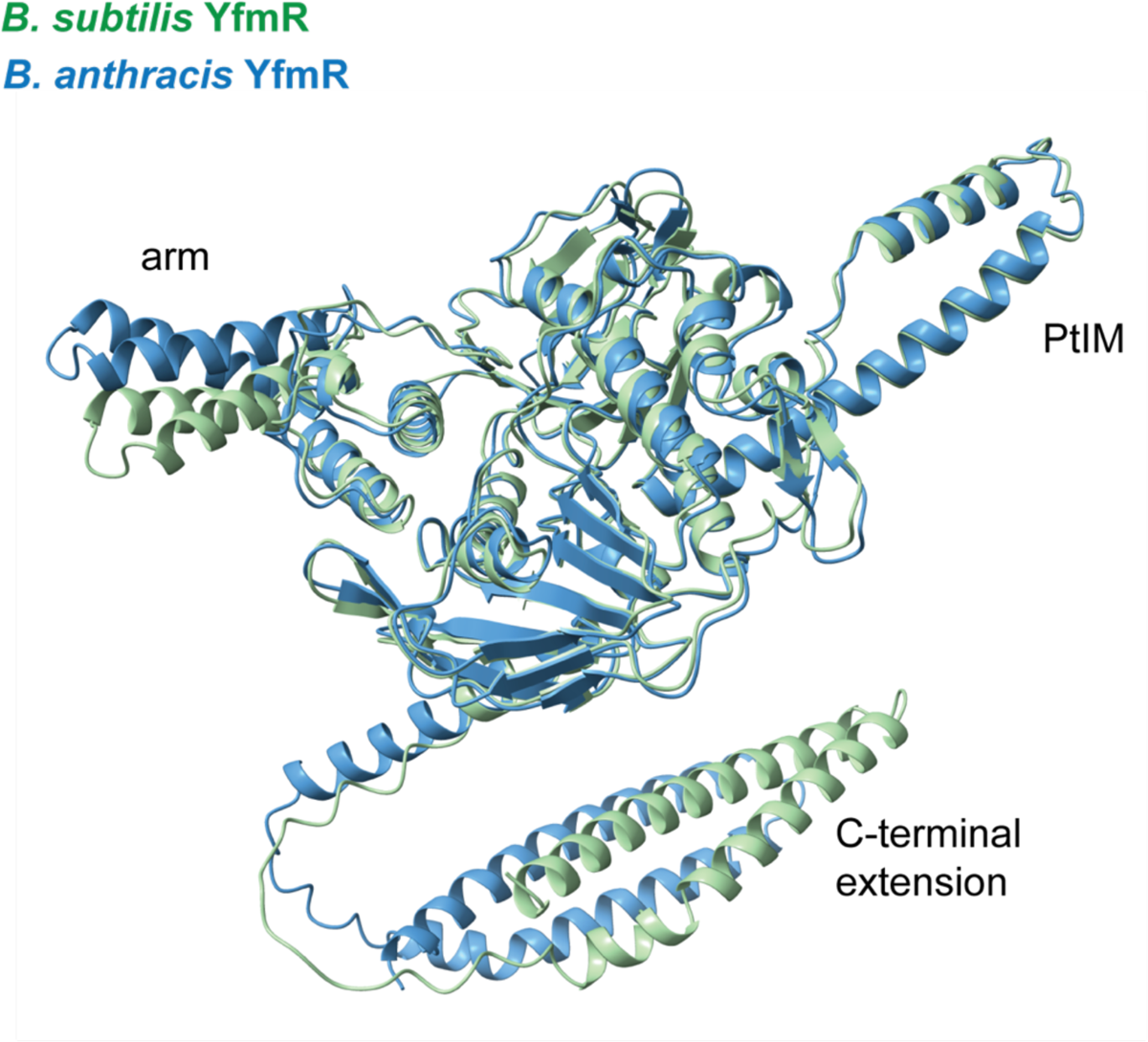
Structural overlay of YfmR from *B. subtilis* and *B. anthracis*. *B. subtilis* and *B. anthracis* YfmR structures were created with AlphaFold and overlayed with UCSF ChimeraX. *B. subtilis* YfmR is displayed in green and *B. anthracis* YfmR is displayed in blue.

**Table S1.**
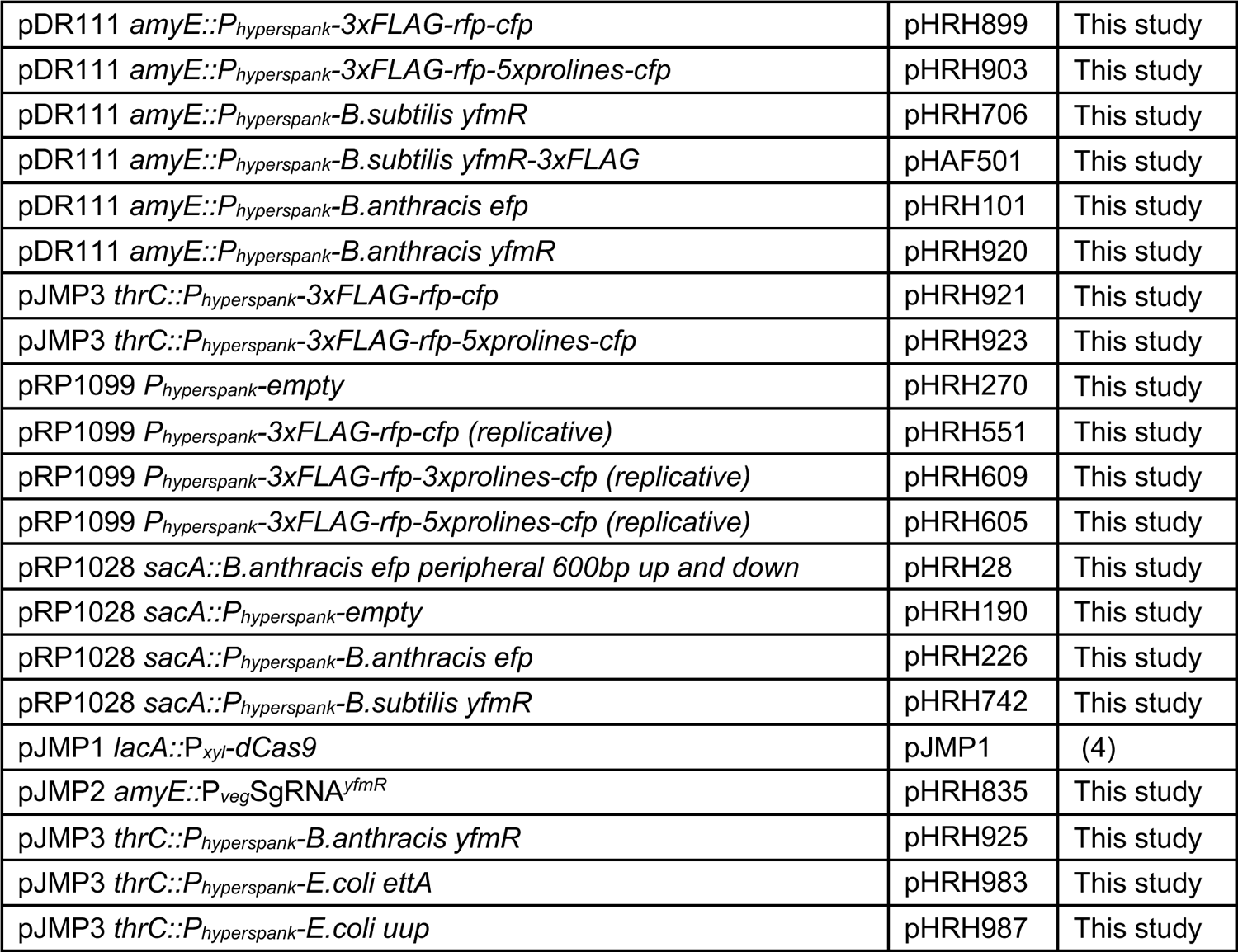
Plasmids.

**Table S2.** Strains.

|  |  |  |
| --- | --- | --- |
| 168 trpC2 *B. subtilis* wild type | HAF1 | (20) |
| 168 trpC2 Δefp::kan | HAF24 | This study |
| 168 trpC2 Δefp | HRH575 | This study |
| 168 trpC2 ΔyfmR::kan | HAF451 | This study |
| 168 trpC2 amyE::P<sub>hyperspank</sub>-*B. subtilis* yfmR-3xFLAG | HAF505 | This study |
| 168 trpC2 Δefp::kan lacA::P<sub>xyl</sub>-dCas9 amyE::P<sub>veg</sub>SgRNA<sup>yfmR</sup> | HRH835 | This study |
| 168 trpC2 amyE::P<sub>hyperspank</sub>-3xFLAG-rfp-cfp | HRH907 | This study |
| 168 trpC2 amyE::P<sub>hyperspank</sub>-3xFLAG-rfp-5xProlines-cfp | HRH908 | This study |
| 168 trpC2 Δefp::kan amyE::P<sub>hyperspank</sub>-3xFLAG-rfp-cfp | HRH909 | This study |
| 168 trpC2 Δefp::kan amyE::P<sub>hyperspank</sub>-3xFLAG-rfp-5xProlines-cfp | HRH910 | This study |
| 168 trpC2 ΔyfmR::kan amyE::P<sub>hyperspank</sub>-3xFLAG-rfp-cfp | HRH911 | This study |
| 168 trpC2 ΔyfmR::kan amyE::P<sub>hyperspank</sub>-3xFLAG-rfp-5xProlines-cfp | HRH912 | This study |
| 168 trpC2 Δefp::kan lacA::P<sub>xyl</sub>-dCas9 amyE::P<sub>veg</sub>SgRNA<sup>yfmR</sup><br>thrC::P<sub>hyperspank</sub>-3xFLAG-rfp-cfp | HRH935 | This study |
| 168 trpC2 Δefp::kan lacA::P<sub>xyl</sub>-dCas9 amyE::P<sub>veg</sub>SgRNA<sup>yfmR</sup><br>thrC::P<sub>hyperspank</sub>-3xFLAG-rfp-5xProlines-cfp | HRH937 | This study |
| 168 trpC2 Δefp::kan lacA::P<sub>xyl</sub>-dCas9 amyE::P<sub>veg</sub>SgRNA<sup>yfmR</sup><br>thrC::P<sub>hyperspank</sub>-*B. anthracis* yfmR | HRH966 | This study |
| 168 trpC2 Δefp::kan lacA::P<sub>xyl</sub>-dCas9 amyE::P<sub>veg</sub>SgRNA<sup>yfmR</sup><br>thrC::P<sub>hyperspank</sub>-*E. coli* etta | HRH1006 | This study |
| 168 trpC2 Δefp::kan lacA::P<sub>xyl</sub>-dCas9 amyE::P<sub>veg</sub>SgRNA<sup>yfmR</sup><br>thrC::P<sub>hyperspank</sub>-*E. coli* uup | HRH998 | This study |
| 34F2 *B. anthracis* Sterne wild type | HAF356 |  |
| 34F2 Δefp | HRH126 | This study |
| 34F2 Δhpf | HRH683 | This study |
| 34F2 Δefp sacA::P<sub>hyperspank</sub>-*B. anthracis* efp | HRH318 | This study |
| 34F2 Δefp sacA::P<sub>hyperspank</sub>-*B. subtilis* yfmR | HRH801 | This study |
| 34F2 pHRH551 P<sub>hyperspank</sub>-3xFLAG-rfp-cfp (replicative) | HRH573 | This study |
| 34F2 pHRH609 P<sub>hyperspank</sub>-3xFLAG-rfp-3xprolines-cfp (replicative) | HRH615 | This study |
| 34F2 pHRH605 P<sub>hyperspank</sub>-3xFLAG-rfp-5xprolines-cfp (replicative) | HRH612 | This study |
| 34F2 Δefp pHRH551 P<sub>hyperspank</sub>-3xFLAG-rfp-cfp (replicative) | HRH572 | This study |
| 34F2 Δefp pHRH609 P<sub>hyperspank</sub>-3xFLAG-rfp-3xprolines-cfp (replicative) | HRH631 | This study |
| 34F2 Δefp pHRH605 P<sub>hyperspank</sub>-3xFLAG-rfp-5xprolines-cfp (replicative) | HRH627 | This study |

**Table S3.**
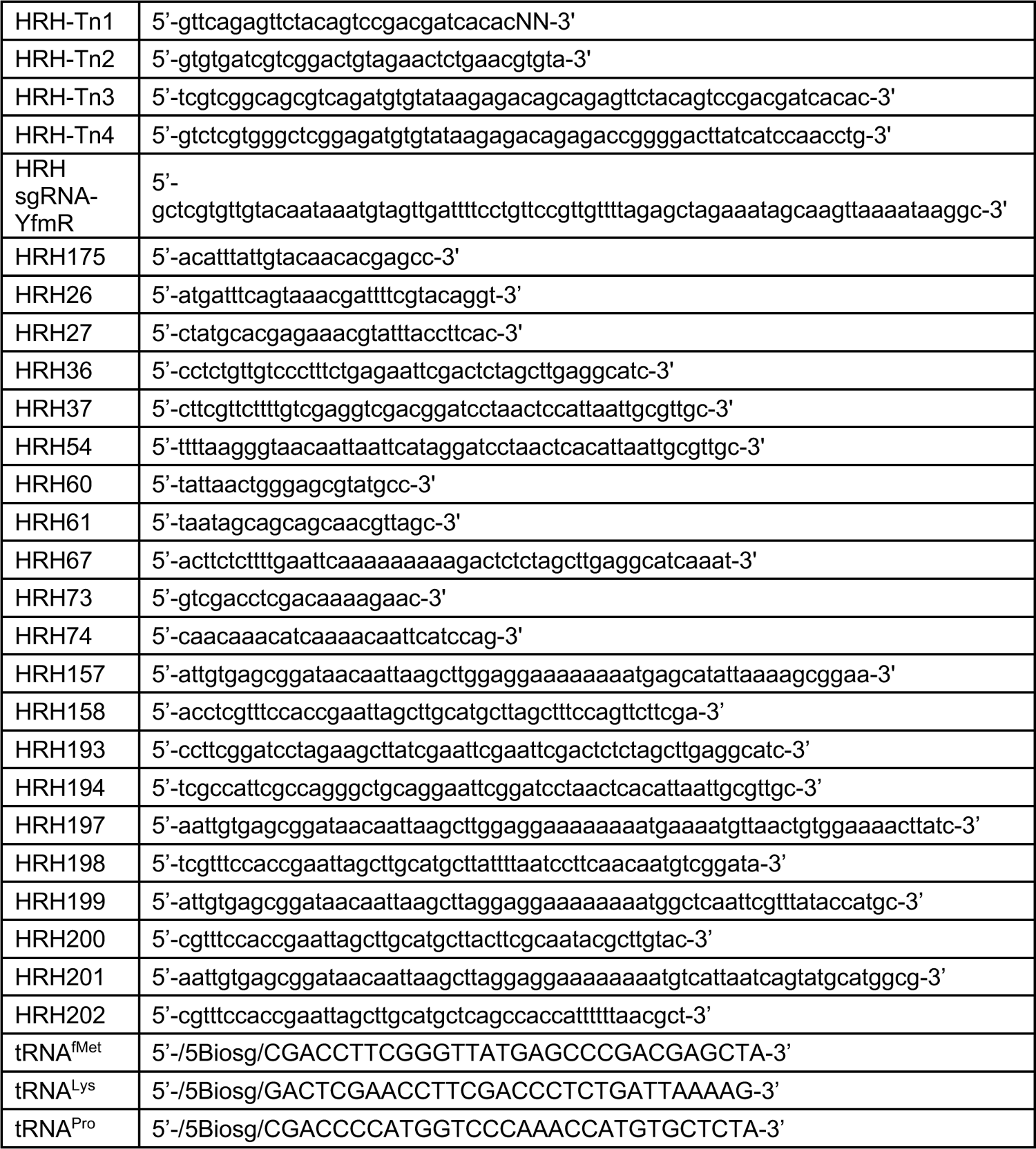
Primers and probes.

**Dataset S1 (separate file).** Excel file containing tallied transposon insertions.

SI References

